# Isoflurane and Surgical Stress Disrupt Fatty Acid and Carbon Metabolism Leading to Cardiomyopathy

**DOI:** 10.1101/2025.10.20.683563

**Authors:** Wendy W Yang, Anna W Chen, Hangnoh Lee, Hui Li, Yun Li, Jin-Gu Lee, Wei-Bin Shen

## Abstract

Aging alters cardiac resilience to anesthetic and surgical stress, yet the molecular basis of these effects remains poorly understood. To define age-dependent cardiac transcriptional responses to isoflurane exposure and operative (ISO/OP) stress, we analyzed gene expression profiles across young adult (3m), late middle-aged (17m), and geriatric mice (27m) following short-term 2 h ISO/OP exposure. At 24 h after cessation, all age groups exhibited distinct cardiac transcriptional signatures separating ISO/OP from sham controls. In young adult hearts, transcriptional alterations 24 hours after cessation of ISO/OP were characterized by dysregulation of small molecule catabolic processes, fatty acid metabolism, disruptions to protein processing in endoplasmic reticulum and cytoskeletal organization. Late middle-aged mice displayed amplified perturbations in lipid metabolism alongside suppression of muscle system and calcium signaling pathways, while old mice showed robust activation of PPAR and AMPK signaling and downregulation of genes governing contractility and morphogenesis. In contrast, geriatric mice showed upregulation of fatty acid metabolic pathways, robust activation of PPAR and AMPK signaling, coupled with suppression of muscle differentiation and actin organization following ISO exposure, indicating a maladaptive metabolic reprogramming. Overlapping DEGs across all age groups converged on pathways regulating oxidative stress, Ca^2+^ handling, hypertrophy, and energy metabolism, suggesting a conserved but age-intensified cardiac stress response. Longitudinal profiling in aged mice revealed persistent transcriptomic remodeling five weeks after stress. Crucially, this remodeling was observed even after ISO exposure alone, indicating that general anesthesia is a primary driver of this long-term effect. This persistent signature was marked by mitochondrial dysfunction and dysregulation of genes associated with diabetic cardiomyopathy, extracellular matrix integrity, and neurodegenerative signaling. Together, these findings identify isoflurane exposure as a potent inducer of persistent, age-dependent metabolic and structural reprogramming in the heart, implicating impaired lipid utilization and mitochondrial homeostasis as central mechanisms linking the perioperative period, and specifically anesthetic exposure, to long-term cardiovascular vulnerability.

## INTRODUCTION

General anesthesia (GA) is an essential element for surgical operations (OP) and diagnostics procedures, but it could also lead to various side effects for the cardiovascular system. Accumulating evidence indicates that these interventions can trigger systemic metabolic and inflammatory responses that extend beyond the perioperative period. While perioperative neurocognitive disorders have received substantial attention [1, 2], the cardiovascular consequences of anesthesia and surgery remain poorly characterized, particularly in the context of aging and noncardiac surgeries that don’t use bypass procedures. However, many studies report significant activation of systemic inflammation, oxidative stress, and neuroendocrine activation following GA/OP [3], which can have secondary effects on cardiac tissue and the cardiovascular system in general. Isoflurane (ISO), a volatile anesthetic widely used in surgical practice, has complex effects on cardiac physiology. In young hearts, ISO is generally well tolerated and can even confer preconditioning benefits against ischemic injury [4, 5]. However, in aged myocardium, anesthetic exposure has been associated with impaired mitochondrial function, altered substrate utilization, and increased oxidative stress [6, 7]. Moreover, a few key studies show that isoflurane (ISO) can have adverse effects on the cardiovascular system, such as induce decreased contraction and worsening oxidative stress in Zucker diabetic fatty rats [8], while a systemic study on occupational exposure to ISO saw dose-dependent effects on cardiac hemodynamics [9]. Despite these observations, how short-term exposure to GA combined with surgical stress (ISO/OP) reprograms cardiac gene expression have not been systematically examined.

Aging is a major risk factor for adverse cardiovascular outcomes in the perioperative setting. Older adults experience increased susceptibility to myocardial injury, arrhythmias, and delayed recovery following anesthesia and surgical stress [10, 11]. This vulnerability stems from progressive declines in cardiac metabolic flexibility, mitochondrial efficiency, and redox homeostasis that limit the heart’s ability to adapt to acute energetic and oxidative stressors. Age-related remodeling of cardiomyocyte bioenergetics includes impaired fatty acid oxidation, altered NAD⁺ metabolism, and increased reliance on glycolytic substrates, which collectively compromise ATP production under stress conditions [12–14]. Concurrently, structural and molecular changes in the extracellular matrix, calcium-handling machinery, and cytoskeletal organization reduce myocardial resilience to hemodynamic fluctuations and inflammatory signaling during surgery. Despite these well-recognized physiological vulnerabilities, the molecular and transcriptional mechanisms through which aging modifies the cardiac response to anesthetic and operative stress remain poorly understood. Elucidating these age-dependent pathways is essential for identifying therapeutic targets to preserve mitochondrial function, stabilize cardiac metabolism, and mitigate postoperative cardiac dysfunction in the growing aging population.

Our previous transcriptomic work demonstrated that ISO/OP exposure in aged mice disrupts neural and behavioral function, producing persistent olfactory deficits, motor frailty, and delayed cognitive decline[15]. These findings suggested that perioperative stress can elicit long-lasting molecular and functional changes within the central nervous system. However, whether similar age-dependent transcriptional remodeling occurs in peripheral organs such as the heart— an organ with high metabolic turnover and sensitivity to oxidative stress—remains unknown. Given the growing recognition of metabolic coupling between the heart and other organ systems [16, 17], understanding how anesthesia and surgery influence cardiac bioenergetics is critical for identifying mechanisms of perioperative vulnerability in aging. In particular, a study using experimental mice models demonstrated that heart failure could lead to significant alterations in hippocampal gene expression and impaired memory function, highlighting the significance of cardiac function to higher cognitive abilities [18].

Despite growing evidence linking age to differential anesthetic sensitivity, little is known about how cardiac transcriptional networks are reprogrammed by anesthetic and surgical exposure across the lifespan. Recent advances in transcriptomic profiling now enable high-resolution characterization of these molecular responses, offering new insights into age-dependent stress adaptations. Here, we investigated the cardiac transcriptional landscape following short-term ISO exposure combined with surgical stress (ISO/OP) in young adult, middle-aged, and old mice. We hypothesized that aging alters the temporal and pathway-specific transcriptional responses to ISO/OP, leading to distinct molecular signatures of metabolic and structural remodeling. By integrating age-stratified transcriptomic analyses, we define dynamic, age-dependent cardiac transcriptional programs that may underlie heightened perioperative vulnerability in the aged heart. We analyzed both acute (24 h) and chronic (5 w) transcriptional responses to determine how aging modulates the heart’s molecular resilience to perioperative stress. Our results reveal that ISO/OP induces rapid, age-dependent dysregulation of fatty acid metabolism, cytoskeletal integrity, and mitochondrial pathways, with persistent remodeling of genes associated with diabetic cardiomyopathy and neurodegenerative signaling in aged mice. These findings identify anesthesia and surgical stress as potent drivers of metabolic and structural reprogramming in the aging heart, providing new insights into the molecular basis of perioperative cardiac vulnerability.

## MATERIALS AND METHODS

### Mouse General Anesthesia and Operation (GA/OP)

All experimental protocols were approved by the Institutional Animal Care and Use Committee (IACUC) at the University of Maryland School of Medicine. Young adult (10–12 weeks, 2.5–3.0-month-old), middle-aged (17-month-old), aged (20-21-month-old), and old (27-month-old) C57BL/6 male mice were sourced from Charles River Laboratories through the National Institute on Aging (NIA). Mice were housed under a 12-hour light/dark cycle with *ad libitum* access to food and water. General anesthesia was induced and maintained with 2% isoflurane in 100% oxygen via a precision vaporizer. Anesthetic depth was monitored by respiratory rate and depth, as well as palpebral and pedal reflexes. Core body temperature was maintained using a heated pad throughout the procedure. After induction, mice were positioned on a surgical station with a nose cone for continuous isoflurane delivery. A midline laparotomy was performed through a 1.5 cm incision extending from the xiphoid process to 0.5 cm above the pubic symphysis, sequentially cutting through the skin, abdominal musculature, and peritoneum. Post-surgery, the incision was treated with 0.25% bupivacaine in sterile saline and closed in layers using 5-0 monofilament nylon sutures. The surgical procedure lasted approximately 10 minutes, after which mice were returned to the anesthesia chamber to complete a 2-hour isoflurane exposure. During recovery, animals were placed on a temperature-controlled pad for 30–60 minutes to stabilize core temperature. Postoperative monitoring included intensive observation for 4 hours following anesthesia/surgery, with daily assessments thereafter. A separate cohort underwent 2-hour isoflurane exposure without surgery, while sham mice received no anesthesia or surgery and remained in their home cages with room air exposure for 2 hours, reflecting clinically relevant conditions. Blinding was not feasible due to the visible abdominal wounds.

For assessment of long-term cardiac tissue, a separate cohort of aged 20-month-old was subjected to laparotomy and 2-hour isoflurane exposure as described in previous publication [15]. In brief, following a battery of neurological behavioral tests that was conducted for up to 38 days, mice were euthanized at 5 weeks after cessation and the hearts were obtained for downstream molecular experiments

### RNA Extraction and Bulk RNA Sequencing (RNAseq)

Following euthanasia at 24 hours and 5 weeks post-ISO/OP, mice were perfused with 50 mL of ice-cold saline. Total RNA was extracted from the heart of both sham and ISO/OP mice using the Direct-zol™ RNA Miniprep Plus kit (Cat# R2073-A, Zymo Research). Extracted RNA samples were sent to Psomagen (Rockville, MD) for mRNA library preparation via Illumina Stranded Total RNA (Ribo-Zero Plus) and sequenced as 150 bp paired-end reads on a NovaSeq X Plus 25B platform (Illumina, CA).

### Statistical and RNAseq Analysis

RNA-Seq reads were aligned to the mouse reference genome (GRCm39) using STAR aligner version 2.7.5 [19], with GENCODE gene annotation (version M33). Gene and isoform expression levels were quantified in transcripts per million (TPM) using RSEM 1.3.3 [20]. Differential expression analysis was conducted using DESeq2 version 1.42.1 with false discovery rates (FDR) calculated via the Benjamini-Hochberg method. Genes with FDR < 0.05 were considered differentially expressed and subjected to downstream pathway enrichment analysis. Gene ontology analysis was performed using clusterProfiler 4.10.1 [21] on the DESeq2 output. Detailed statistical analyses for each assay are provided in the figure legends, with a significance threshold set at p ≤ 0.05.

## RESULTS

### Short-term ISO/OP exposure in young adult mice dysregulates cardiac fatty acid metabolism and cytoskeletal gene expression after 24 h of cessation

To evaluate the effects that short-term general anesthesia would have on the cardiac system of young adult mice, we subjected 3-mon-old (20-30 human years) C57BL/6 male mice to laparotomy and 2h ISO exposure. Transcriptomic profiling of the heart at 24h after cessation revealed clear separation between sham and Iso/OP groups in PLSDA analysis (Fig. 1A). Following pairwise comparison, DEGs depicted in the volcano plot showed an equal number of up- and down-regulated genes in the heart of Iso/OP mice (Fig. 1B), with the top ten DEGs consisting of four upregulated (*Serpina3n, Mid1ip1, Lcn2* and *Tef*) genes, along with six downregulated genes (*Eln, Col5a3, Arntl, Npas2, Slc41a3, Adam19*). Pathway enrichment analysis of the upregulated DEGs demonstrated that “small molecule catabolic process”, “fatty acid metabolic process”, and “carboxylic acid catabolic process” are the top 3 molecular processes in GO terminology (Fig. 1C). Whereas KEGG pathways revealed “protein processing in ER”, “circadian rhythm” and “fatty acid degradation” as the top affected pathways (Fig. 1D). Enrichment analysis of the downregulated DEGs shows that “cell-substrate adhesion”, “actin filament organization” and “extracellular matrix organization” are the main processes being affected in GO terminology (Fig. 1E). Moreover, further analysis revealed that “focal adhesion”, “salmonella infection”, and “cytoskeleton in muscle cells” are the most significantly enriched pathways in KEGG (Fig. 1F). Representative genes from the top three enriched GO processes were visualized as heatmaps for up- and down-regulated DEGs (Fig. 1G–H). To further illustrate pathway-specific expression changes, normalized transcript abundance (TPM) of key DEGs within each enriched process was quantified and presented as bar graphs, accounting for library size and sequencing depth (Fig. 1I).

**Figure 1.**
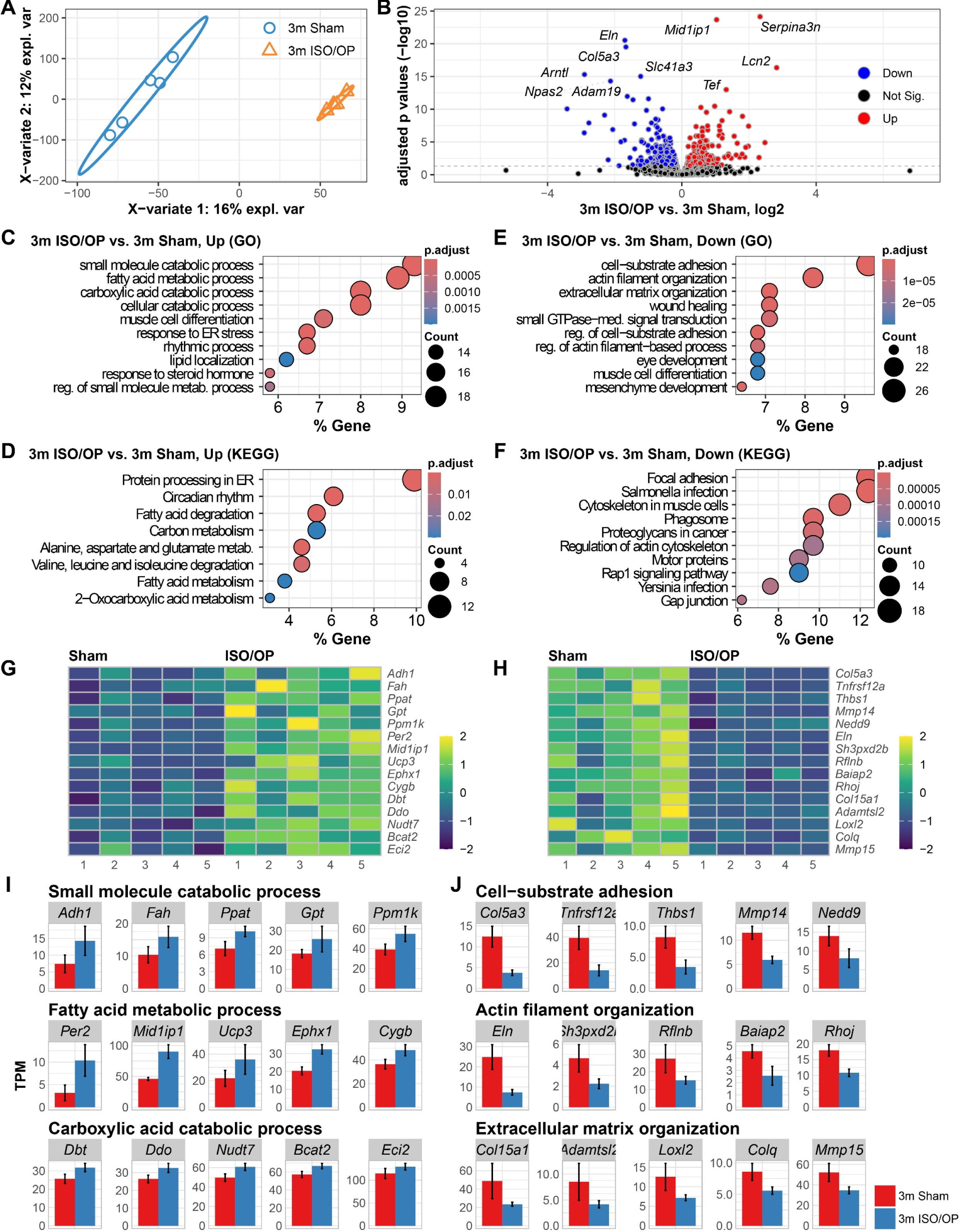
ISO/OP drives acute alterations to transcriptomic profile of cardiac tissue in young adult mice at 24h after isoflurane (ISO) exposure. (**A**) PLS-DA plot for all normalized transcriptome genes is depicted for young adult 3-month-old (3m) mice with or without ISO exposure following laparotomy. (**B**) Volcano plot displaying differentially expressed genes (DEGs) in the 3m ISO/OP vs. 3m Sham comparison. (**C-D**) Pathway enrichment analysis of upregulated DEGs with Gene Ontology molecular processes (**C**) and KEGG pathways (**D**). (**E-F**) Pathway enrichment analysis of downregulated DEGs with Gene Ontology molecular processes (**E**) and KEGG pathways (**F**). Z-score normalization was applied to gene expression values. Heatmaps illustrate sample-wise variation (columns) across genes (rows). Color intensity reflects relative transcript abundance across samples: warmer colors for higher, cooler for lower. (**G-J**) Heatmap and bar graph of genes involved in the top three enriched pathways. n = 5 mice/group.

### Middle-aged mice exhibit impaired fatty acid metabolism and muscle system processes 24 h after cessation

We next examined the effects that Iso/OP may have on 17mon old mice, which are the equivalent of 51–54 years of age for human, a late middle-aged to aged phase. Transcriptomic data at 24h after cessation showed that Iso/OP contributed to 27% of the variation observed between groups (Fig. 2A). Whereas pairwise comparison showed a significant number of total DEGs (Fig. 2B), the most notable ones consisted of four upregulated genes (*Pdk4, Serpine1, Mt2, Serpina3n*), along with two downregulated ones (*Myl7, Fgf12*). Pathway enrichment analysis with GO terms demonstrated heightened activity in “amide metabolic process”, “fatty acid metabolic process”, and “alcohol metabolic process” (Fig. 2C). In KEGG terms, we found a coordinated upregulation of genes related to “cytokine-cytokine receptor interaction”, “protein processing in ER”, and “lipid and atherosclerosis” processes (Fig. 2D). In contrast, downregulated DEGs were found to encompass “muscle system process”, “striated muscle cell differentiation” and “regulation of metal ion transport” in GO molecular processes, suggesting that ISO/OP were affecting muscle cells in the heart at 24h after cessation (Fig. 2E). Moreover, KEGG pathways revealed that genes involved in “cytoskeleton in muscle cells”, “calcium signaling pathway” and “cardiac muscle contraction” were being downregulated (Fig. 2F). To visualize the expression patterns driving the GO biological processes, we generated heatmaps of the key genes belonging to these pathways (Fig. 2G-H), which normalized transcript abundance (TPM) of representative DEGs within each GO category was quantified and visualized as bar graphs, adjusted for library size and sequencing depth (Fig. 2I).

**Figure 2.**
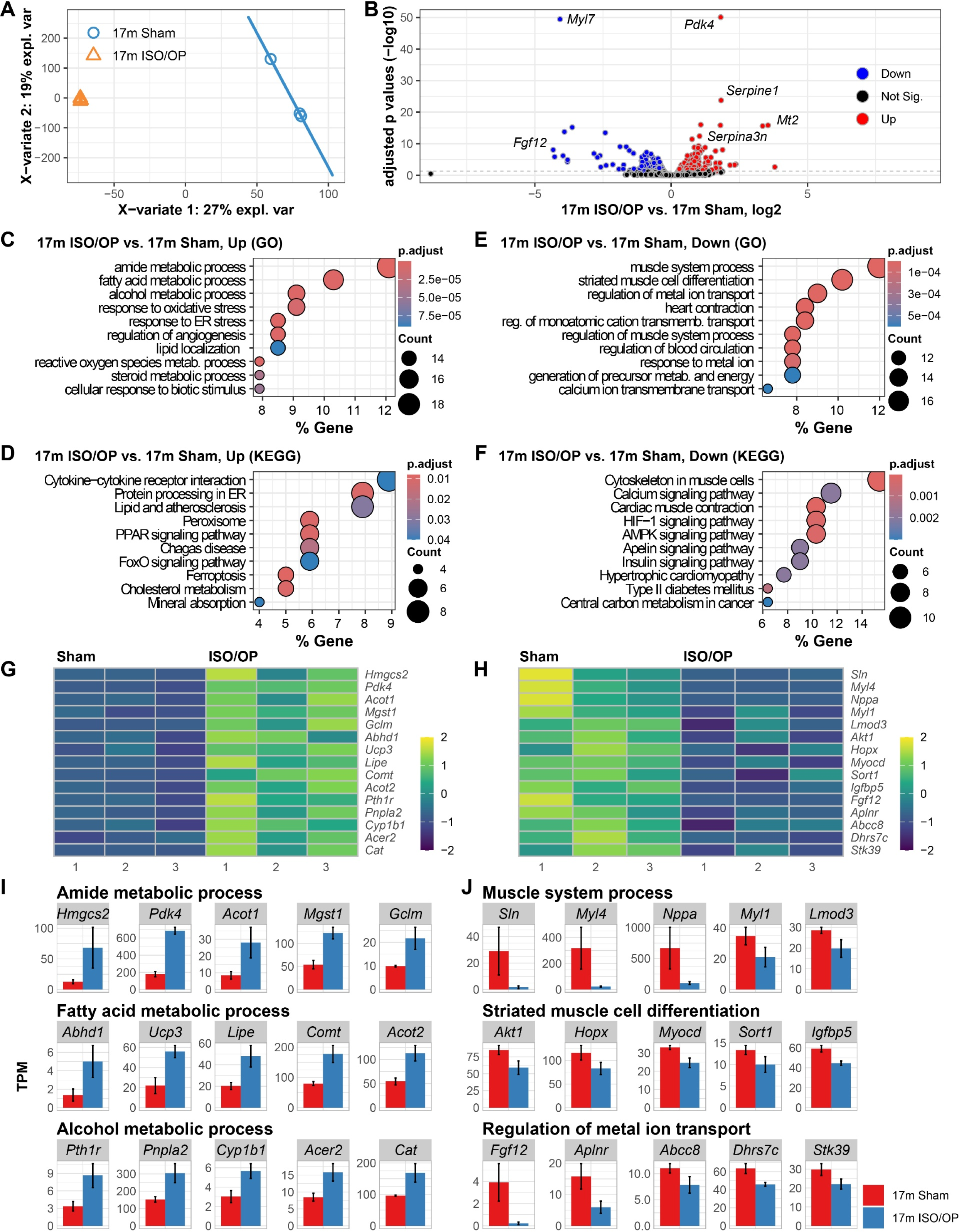
Heart transcriptomic profiling at 24 h after short-term ISO/OP reveals altered metabolic and muscle cell pathways in aged 17-month-old (17m) mice. (**A**) PLS-DA showing clear separation of heart transcriptomes across groups. (**B**) Volcano plot of differentially expressed genes (DEGs) comparing 17m ISO/OP vs. 17m Sham. (**C-D**) Enriched GO molecular processes (**C**) and KEGG pathways (**D**) for upregulated DEGs. (**E-F**) Enriched GO molecular processes (**E**) and KEGG pathways (**F**) for downregulated DEGs. Heatmaps of representative genes contributing to the top enriched GO term pathways (**G, H**). (**I–J**) Bar plots illustrating expression of selected DEGs linked to the top three enriched pathways in up-(**I**) and down-(**J**) regulated DEGs. n=3 mice/group.

### Old mice exposed to ISO exhibit upregulation of fatty acid metabolic processes and downregulation of muscle cell differentiation

After exploring the effects of ISO/OP on the cardiovascular system of young and late middle-aged, we examined the effects this procedure may have on old 27-month-old mice, which are equivalent to 80 years of age in humans. PLS-DA of all normalized genes at 24h after cessation showed a clear separation between sham and ISO/OP mice, with the procedure contributing to 17% of the variation between groups (Fig. 3A). In addition, DEGs between groups showed a robust change in the cardiac transcriptomic profile (Fig. 3B), with notable upregulation observed in *Slc27a1, Pnpla2, Lrg1, Rhobtb1 and Per1*, which span core aspects of cardiac physiology, including energy metabolism (*Slc27a1, Pnpla2*), fibrosis regulation (*Lrg1*), vascular tone maintenance (*Rhobtb1*), and hemodynamic circadian modulation (*Per1*). In addition, the three notable downregulated genes, *Tfrc*, *Tuba8* and *Wdr1* are reported to be involved in iron metabolism, cytoskeletal dynamics, and actin filament regulation, which are critical to cardiac cell physiology. To extract biological meaning from the list of regulated genes, we performed a functional enrichment analysis using GO molecular process and KEGG pathway databases. Pathway enrichment analysis of the upregulated DEGs showed “small molecule catabolic”, “cellular catabolic” and “lipid catabolic” as the top GO term processes being mediated by the ISO/OP treatment (Fig. 3C).

**Figure 3.**
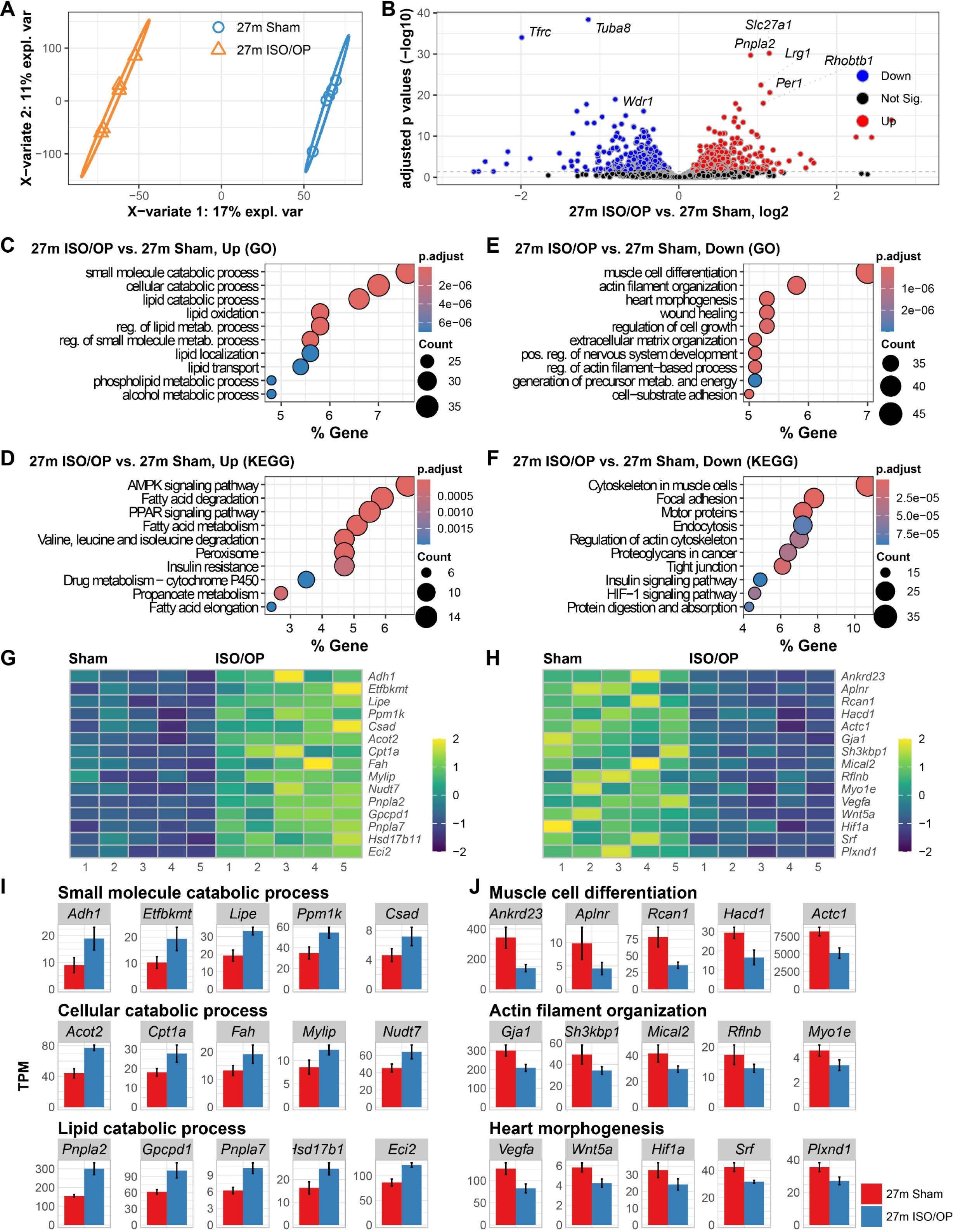
Short-term isoflurane (ISO) exposure alters acute transcriptomic responses in the heart of aged 27-month-old (27m) mice 24 h after cessation. (**A**) PLS-DA of heart transcriptomes across groups shows clear separation. (**B**) Volcano plot of differentially expressed genes (DEGs) comparing 27m ISO/OP vs. 27m Sham. (**C-D**) Enriched GO molecular processes (**C**) and KEGG pathways (**D**) for upregulated DEGs. (**E-F**) Enriched GO molecular processes (**E**) and KEGG pathways (**F**) for downregulated DEGs. Heatmaps of representative genes contributing to the top enriched GO term pathways (**G, H**). (**I–J**) Bar plots illustrating expression of selected DEGs linked to the top three enriched pathways in up-(**I**) and down-(**J**) regulated DEGs. n=5 mice/group.

Whereas in KEGG analysis, “AMPK signaling pathway”, “Fatty acid degradation”, and “PPAR signaling pathway” are the top enriched pathways (Fig. 3D). In contrast, pathways involved in “muscle cell differentiation”, “actin filament organization” and “heart morphogenesis” were highly prominent in the downregulated DEGs (Fig. 3E). While KEGG analysis revealed a strong signature in genes involved with “cytoskeleton in muscle cells”, “focal adhesion” and “motor proteins” (Fig. 3F). Finally, we visualized the expression patterns of key genes from the enriched GO processes using heatmaps (Fig. 3G-H), and we quantified the normalized transcript abundance (TPM) of representative DEGs from each category using bar graphs, adjusted for library size and sequencing depth (Fig. 3I).

### The acute cardiac transcriptional response to ISO/OP is age-dependent after 2 h of exposure

We next explored whether key ISO/OP mediated targets could be isolated from across different age groups by examination of overlapping genes. Amongst the upregulated DEGs derived from 3m, 17m and 27m mice, a total of 19 genes showed overlap in ISO/OP vs. sham mice comparison groups (Fig. 4A). The expression pattern of these key genes across the three different age groups were visualized as a heatmap following normalization by their respective sham groups (Fig. 4B). Key notable genes in this list are *Cyp4b1*, which has been reported to be upregulated in a model of thyroid-stimulating hormone induced cardiac hypertrophy *and Ephx1,* which has been reported to impair cardiac recovery after ischemia [22, 23]. In addition, the gene *Fbxo31* has been reported to mediate senescence in endothelial cells, which is a leading mechanism for atherosclerosis [24]. Moreover, increased expression levels of *Lcn2* have been reported to promote cardiomyocyte apoptosis and inflammation [25, 26]. Overall, these results suggest that acute ISO/OP in aged mice could induce an upregulation of genes related to oxidative stress, Ca^2+^ handling, hypertrophy regulation, and lipid metabolism, which seem to increase with age. Next, we examined the overlapping genes derived from downregulated DEGs, which yielded a total of 24 genes (Fig. 4C), which are visualized as heatmaps to explore their changes across the three different age groups (Fig. 4D). Key genes encompassed in these processes include *Akt1, Asph, Mtr, Tfrc, Pcdh12* and *Ppp1r3c* which have been identified to play distinctive roles in cardiac function by regulating energy homeostasis, calcium dynamics and metabolism. Moreover, the genes *Adam19, Akt1, Arntl* and *Cd93* have been reported to modulate cellular senescence and cardiac aging [27–30]. Together, these findings highlight that ISO/OP exposure elicits convergent transcriptional responses across ages, with fatty acid metabolism and energy regulatory pathways, along with dysregulation to genes involved with cellular senescence emerging as core molecular signatures of cardiac stress at 24 h post-exposure.

**Figure 4.**
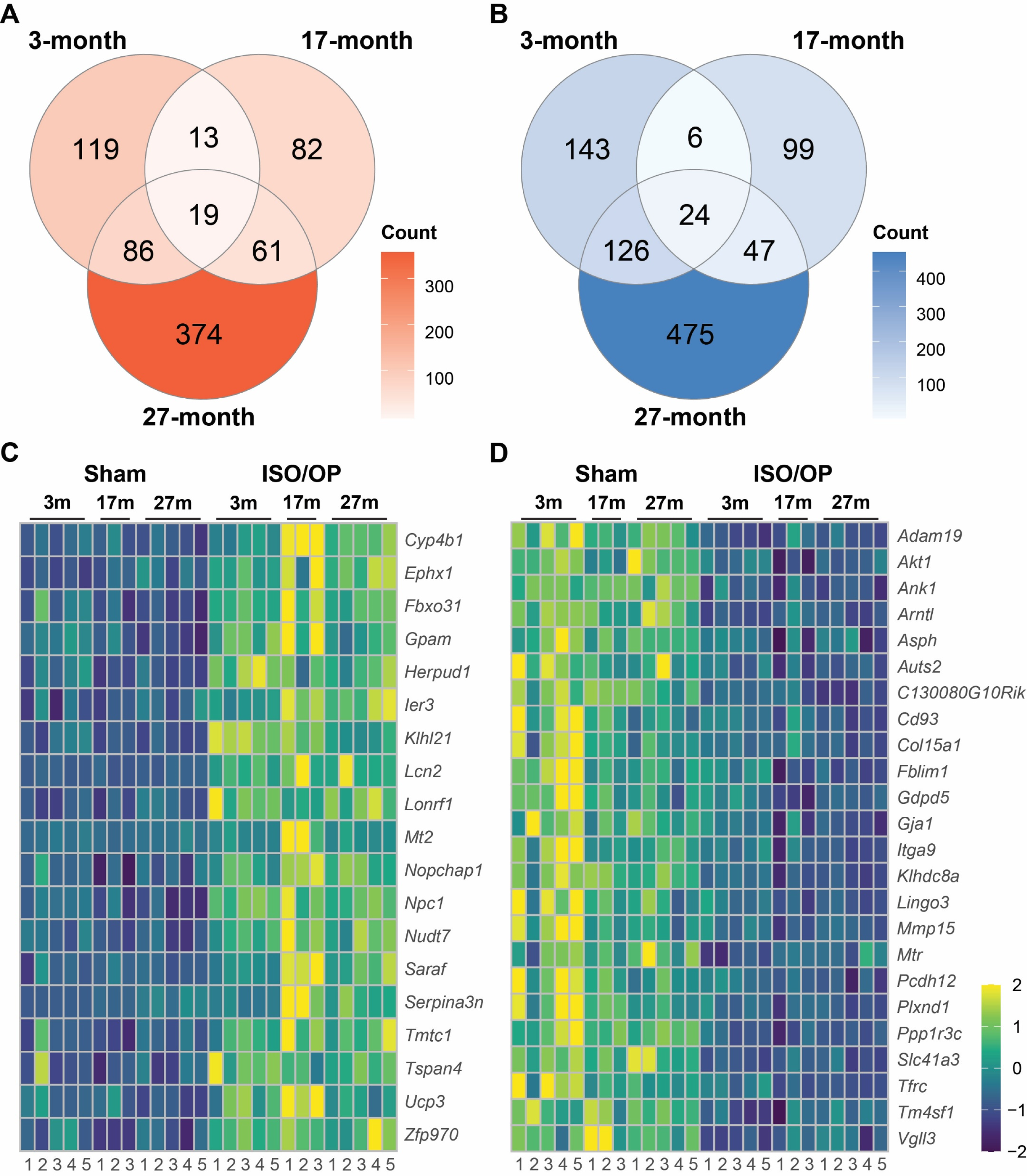
Overlap analysis of differentially expressed genes (DEGs) across different age groups identify key genes. (**A-B**) Venn diagrams of the up-(**A**) and down-(**B**) regulated DEGs obtained from ISO/OP vs. Sham comparison across different age groups. (**C-D**) Heatmaps of overlapping genes in up-(**C**) and down-(**D**) regulated gene sets. n=3-5 mice/group

### ISO/OP treatment induces long-term dysregulation of diabetic cardiomyopathy–associated genes in aged mice

A previous study from our group demonstrated that ISO/OP exposure in 20-month-old mice induces persistent olfactory dysfunction, reduced limb strength, and impaired motor coordination indicative of frailty, accompanied by delayed cognitive deficits and increased apathy in the same cohort [15]. Emerging evidence and scientific statements from the American Heart Association have shown unequivocally that the brain and heart are interdependent organ systems with intricate pathogenic mechanisms that link heart conditions with microstructural changes in the brain [16–18]. Building on these findings, we subsequently collected heart tissues from the same aged mice at 5 w after cessation to investigate the long-term cardiovascular effects of ISO/OP. RNA sequencing was performed on chronic heart samples collected 5 weeks after surgery from aged mice that had undergone either sham procedures or 2 h of ISO/OP. Initial analysis of the cardiac transcriptional profiles showed clear separation between Sham and ISO/OP mice in PLSDA, with the surgery contributing to 19% of the total variance (Fig. 5A). Pairwise comparisons further identified distinct molecular alterations, with genes such as *Lars2* and *Atf3* showing marked downregulation, while *mt-Nd3, Ppm1k, Mylk4, mt-Atp8,* and *mt-Co2* were significantly upregulated in ISO/OP-exposed mice (Fig. 5B). To interpret the biological implications of the upregulated DEGs, we performed Gene Ontology (GO) enrichment analysis. This revealed a strong signature of genes involved in ‘RNA splicing’, ‘membraneless organelle assembly’, and ‘regulation of protein-containing complex assembly’ (Fig. 5C). Consistent with these findings, KEGG pathway analysis highlighted a significant enrichment for pathways associated with neurodegeneration, including ‘amyotrophic lateral sclerosis’, ‘prion disease’,‘diabetic cardiomyopathy’ and ‘proteoglycans in cancer’ (Fig. 5D). Consecutively, the top downregulated pathways consisted of “generation of precursor metabolism and energy”, “muscle cell differentiation”, and “mitochondrion organization” in GO terms (Fig. 5E), while KEGG analysis also showed a strong signature in neurodegeneration, with Alzheimer and Parkinson diseases as the next most enriched pathways (Fig. 5F). To further illustrate these transcriptional shifts, we visualized the expression patterns of key genes within enriched GO categories as heatmaps (Fig. 5G–H) and quantified representative DEGs from each pathway as normalized TPM bar plots adjusted for library size and sequencing depth (Fig. 5I).

**Figure 5.**
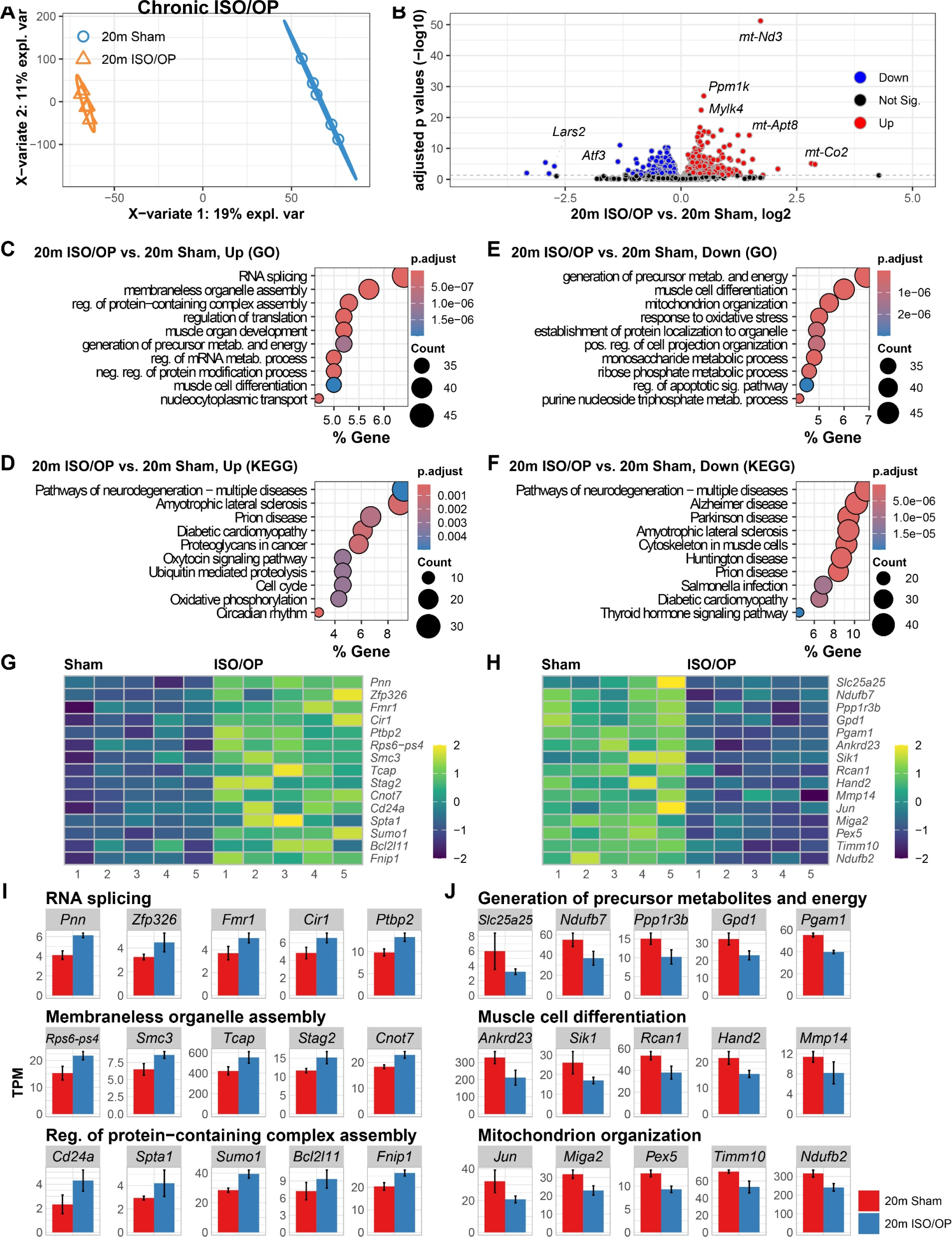
Chronic cardiac transcriptomes at 5 weeks post-cessation reveal persistent alterations to cellular assembly and energy homeostasis in aged mice. (**A**) Multivariate analysis (PLS-DA) reveals persistent divergence of cardiac transcriptomes induced by laparotomy and isoflurane exposure (ISO/OP). (**B**) Volcano plot of differentially expressed genes (DEGs) comparing 20m ISO/OP vs. 20m Sham. (**C-D**) Enriched GO molecular processes (**C**) and KEGG pathways (**D**) for upregulated DEGs. (**E-F**) Enriched GO molecular processes (**E**) and KEGG pathways (**F**) for downregulated DEGs. Heatmaps of representative genes contributing to the top enriched GO term pathways (**G, H**). (**I–J**) Bar plots illustrating expression of selected DEGs linked to the top three enriched pathways in up-(**I**) and down-(**J**) regulated DEGs. n=5 mice/group.

In the KEGG pathway analysis for upregulated DEGs, much of the top three enrichments were related to neurodegeneration and neuroinflammation, but the fourth term was diabetic cardiomyopathy (CM), which was specific to cardiac tissue and needed further interrogation. Notable genes that mediate this pathological process include *Rac1, Ppp1cb, Ndufs4, Atp5j, Mpc1*, along with the mitochondrial DNA genes *mt-Nd2, mt-Co2, mt-Atp8, mt-Nd4 and mt-Cytb*, which were upregulated in ISO/OP mice (Fig. 6A). In contrast, downregulated DEGs that overlap with diabetic cardiomyopathy included *Col1a1, Atp5d, Ctsd, Ndufs7, Ndufb9, Ppara, Col3a1, Slc25a4, Mmp2* and *Cox8a* (Fig. 6B). For rigorous quantification, gene-level expression was depicted as bar graphs with normalized TPM plotted on the y-axis (Fig. 6C). Collectively, these findings reveal that ISO/OP induces persistent, transcriptome-level remodeling of the aged heart, marked by mitochondrial and metabolic dysregulation consistent with diabetic cardiomyopathy and neurodegenerative signaling pathways.

**Figure 6.**
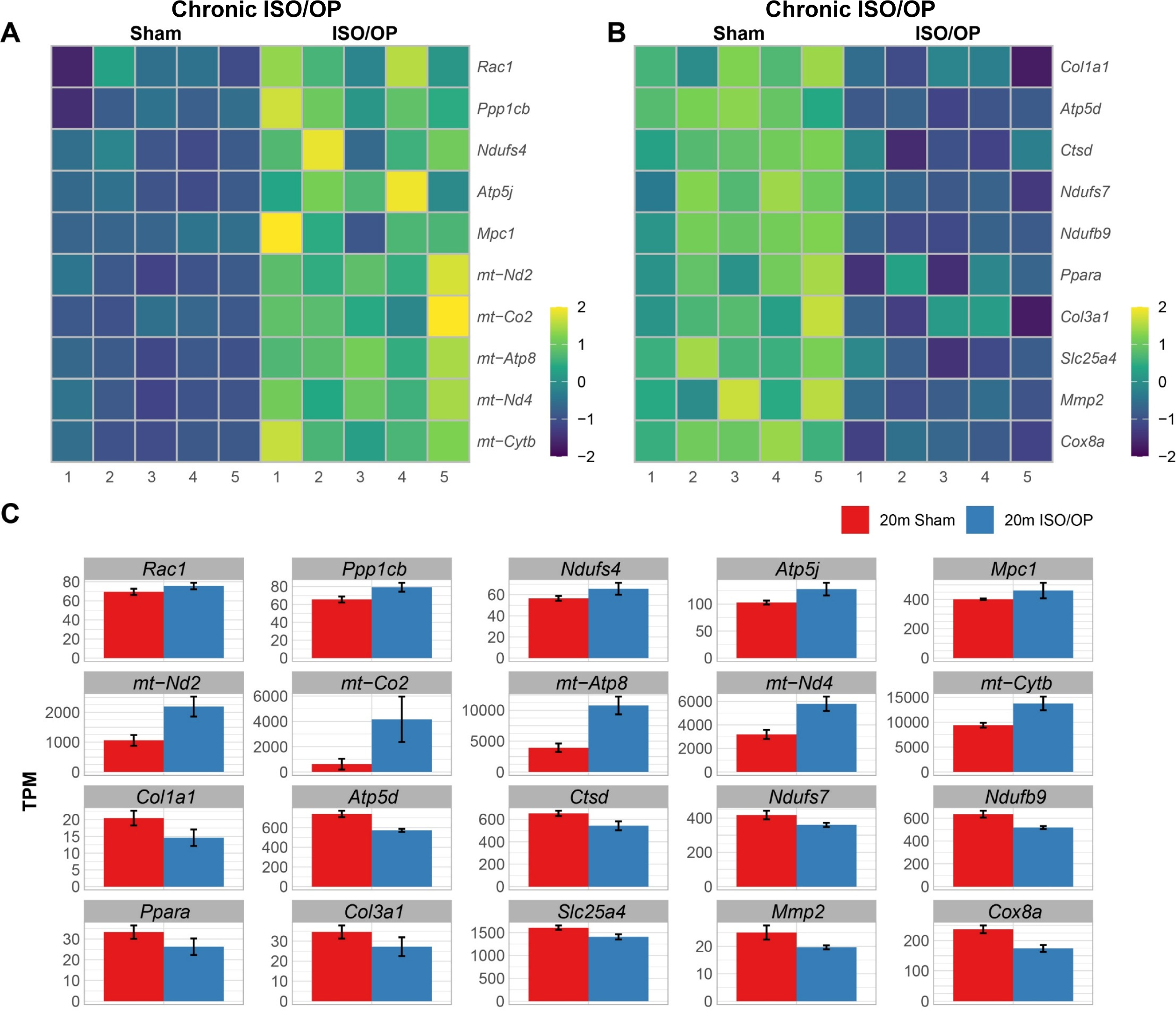
Chronic cardiac transcriptomes at 5 weeks post-cessation show alterations to diabetic cardiomyopathy genes in aged mice. (**A-B**) Heatmaps depict expression patterns of the up-(**A**) and down-(**B**) regulated genes involved with cardiomyopathy, which illustrates sample-wise variation (columns) across genes (rows). Z-score normalization was applied to gene expression values. Color intensity reflects relative transcript abundance across samples, with warmer colors for higher, cooler for lower. Heatmap and bar graph of genes involved in the top three enriched pathways. (**C-D**) Bar graph of the genes related to cardiomyopathy. n=5 mice/group.

In order to explore whether the molecular changes observed were attributable to ISO exposure or surgical stress, we next examined the chronic heart samples obtained from ISO alone groups in the same aged 20m study, in which we also observed impairments in brain and neuromuscular function [15]. PLSDA of the cardiac transcriptomes revealed a distinct separation between sham and ISO mice at 5w after cessation, with 17% of the total variance being found attributable to GA exposure (Fig. 7A). Pairwise comparisons further identified distinct molecular alterations, with genes such as *Actg1*, *Midn*, *Slc41a3*, *Col1a1*, *C1qc* and *Smim7* showing marked downregulation, while *B3galt2* and *Pcmtd1* were significantly upregulated in 20m mice that were exposed to ISO for 2h (Fig. 7B). Afterwards, enrichment analysis of the upregulated DEGs was employed to identify GO terms that were over-represented. GO terminology showed significant enrichment in genes that were related to “eye development”, “stem cell population maintenance”, and “maintenance of cell number” processes (Fig. 7C). On the other hand, downregulated DEGs were shown to be enriched in genes that mediate “cellular response to TGF-beta stimulus”, “response to transforming growth factor beta”, and “connective tissue development” (Fig. 7D). Complementing this, KEGG pathway analysis demonstrated significant enrichment in “cytoskeleton in muscle cells”, “focal adhesion”, and “diabetic cardiomyopathy” pathways amongst the DEGs that were downregulated by ISO exposure (Fig. 7E). Key genes derived from the top three up- and down-regulated pathways were visualized as heatmap to further explore their expression patterns within chronic heart samples (Fig. 7F-G), while quantification of the normalized TPM values were represented as bar graphs for each gene (Fig. 7H-I).

**Figure 7.**
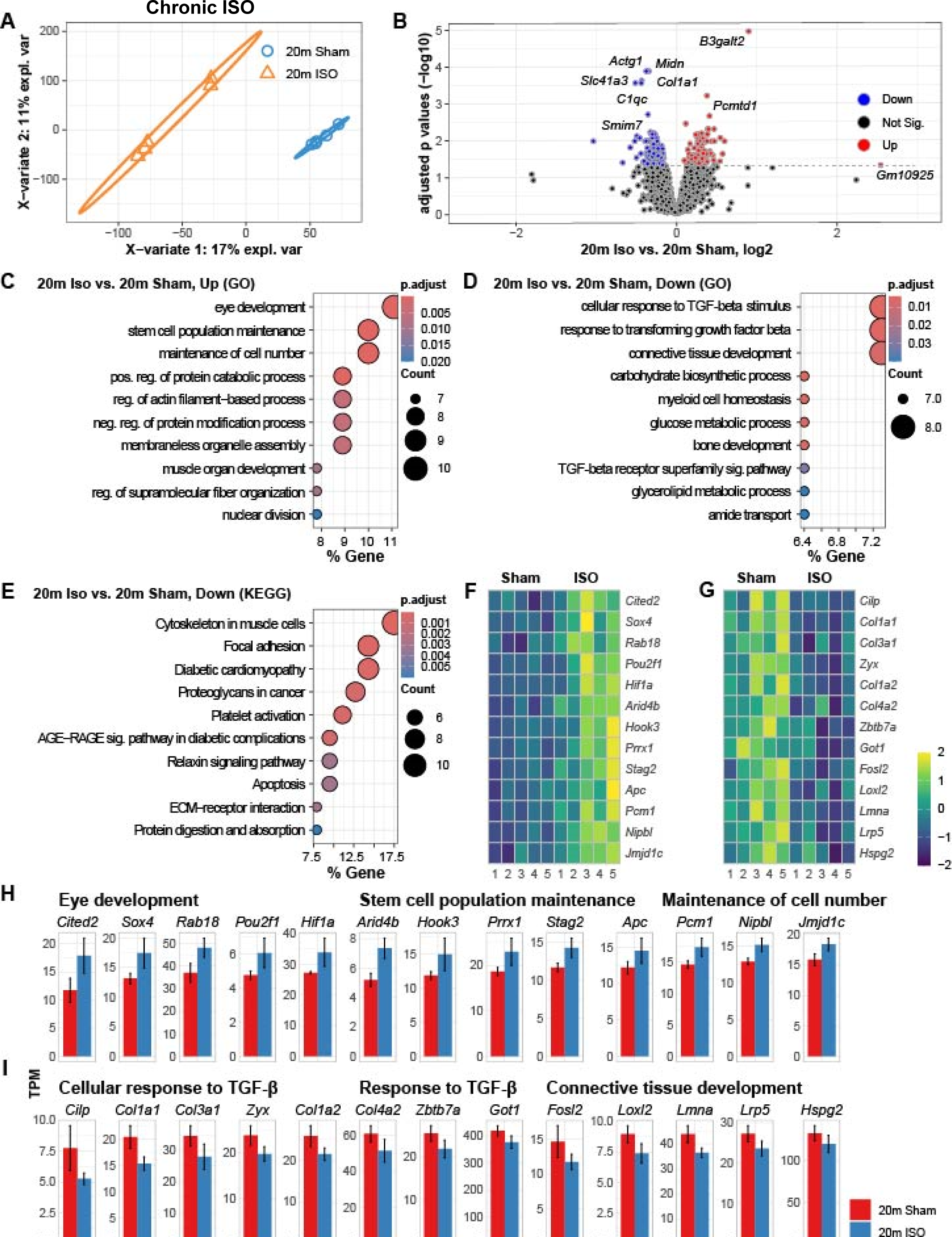
RNA sequencing at 5 weeks after exposure to Isoflurane (ISO) alone reveals persistent alterations to the cardiac transcriptomic profile of aged mice. (**A**) PLS-DA shows clear divergence of cardiac transcriptomes induced by 2 h isoflurane exposure (ISO). (**B**) Volcano plot of differentially expressed genes (DEGs) comparing 20m ISO vs. 20m Sham. (**C-D**) Enriched GO molecular processes for up-(**C**) and down-(**D**) regulated DEGs. (**E**) Enriched KEGG pathways for downregulated DEGs. (**F-G**) Heatmaps of representative genes contributing to the top enriched GO term pathways. (**H–I**) Bar plots illustrating expression of selected DEGs linked to the top three enriched pathways in up-(**I**) and down-(**J**) regulated DEGs. n=5 mice/group.

## DISCUSSION

In the present study, our findings demonstrate that even short-term exposure to GA by inhaled ISO exposure and laparotomy can elicit pronounced, age-dependent transcriptional reprogramming in the heart, highlighting the cardiac system’s unexpected sensitivity to perioperative challenges. In young adult mice, ISO/OP induced acute dysregulation of fatty acid metabolism and cytoskeletal organization, indicating an early metabolic inflexibility and structural remodeling response. With advancing age, these molecular perturbations became more pronounced and diversified: aged 17m mice showed suppression of muscle system processes and calcium signaling pathways, while aged mice exhibited robust activation of lipid catabolic, PPAR, and AMPK signaling pathways coupled with downregulation of genes involved in cardiac morphogenesis and contractile function. The convergence of upregulated fatty acid metabolic and oxidative stress–related genes across all age groups suggests that metabolic stress is a core component of the acute cardiac response to ISO/OP. Importantly, chronic analysis revealed that aged mice failed to fully recover, displaying persistent transcriptomic remodeling consistent with mitochondrial dysfunction and diabetic cardiomyopathy signatures, even those that were exposed to 2 h ISO alone without surgery. Together, these findings identify ISO as a driver of metabolic and structural cardiac stress that escalates with age, offering a potential mechanistic link between perioperative exposure, impaired energy homeostasis, and the heightened cardiovascular vulnerability observed in older patients.

In young adult hearts, ISO/OP exposure triggered acute dysregulation of genes involved in fatty acid metabolism, ER-associated protein processing, and cytoskeletal organization. Because the myocardium derives over 70 % of its ATP from fatty acid oxidation (FAO) under physiological conditions [31], even transient suppression of FAO genes suggests early energetic stress. Enrichment of ER protein-processing transcripts indicates that perioperative exposure imposes a temporary proteostatic load, consistent with prior evidence that volatile anesthetics such as isoflurane disrupt mitochondrial function and activate ER stress pathways in the CNS [32–34]. In addition, studies also show that isoflurane can destabilize the actin cytoskeleton and weaken integrin-mediated adhesion [35–37], which may reflect a broader cellular vulnerability to structural stress. While our RNA-seq data cannot resolve these mechanisms directly, the transcriptional profile observed here aligns with a pattern of transient metabolic and proteostatic challenge, emphasizing that even brief anesthesia and surgical stress can perturb cardiomyocyte homeostasis.

At 24 h post-ISO/OP, converging DEGs across ages indicate disrupted fatty acid metabolism and energy regulation, consistent with age-related declines in mitochondrial β-oxidation and lipid accumulation [12, 14]. Isoflurane also induces ER stress with CHOP and caspase-12 activation and AMPK-driven autophagy [32, 38, 39]. Although pathway enrichment was not significant, several DEGs are known to participate in cardiac aging processes. Aging further amplifies metabolic imbalance through impaired mitophagy, reduced chaperone function, and baseline SASP accumulation [14, 40]. Additionally, ISO-induced cytoskeletal disruption via p75NTR–RhoA impairs mechanotransduction and homeostasis [41]. Together, ISO/OP provokes a multifaceted stress response—metabolic dysfunction, ER stress, and senescence-related gene activation—exacerbated in aged myocardium with diminished compensatory capacity.

At 5w post-ISO/OP, the cardiac transcriptome shifts from acute stress responses to chronic structural and metabolic remodeling. Upregulation of RNA splicing and processing genes suggests ongoing transcriptional reprogramming typical of heart failure, where altered splicing of sarcomeric and calcium-handling proteins impairs contractility and metabolism [42, 43]. Concurrent enrichment of diabetic cardiomyopathy-like signatures points to sustained metabolic stress and disrupted mitochondrial energetics, while the downregulation of anabolic and mitochondrial organization pathways reflects declining bioenergetic efficiency[44]. Overall, our findings suggest that a single short-term exposure to ISO/OP can drive maladaptive remodeling marked by aberrant splicing, mitochondrial disarray, and hypometabolic degeneration akin to late-stage heart failure long after cessation.

Finally, by isolating chronic-timepoint DEGs related to diabetic CM, we found many upregulated mitochondrial genes associated with hyperglycemia-driven dysfunction. ISO exposure induces transient hyperglycemia by impairing insulin signaling and glucose uptake, as shown in rodent, canine, and human studies [45–48]. Chronic hyperglycemia also triggers ER stress—a hallmark of diabetic CM—by overwhelming protein-folding capacity and activating maladaptive UPR signaling [49]. Prolonged ER stress disrupts calcium balance, elevates oxidative stress, and promotes mitochondrial dysfunction and apoptosis [50–52]. Together with dysregulated lipid catabolism observed acutely, these data suggest ER stress as a key mediator linking ISO-induced metabolic overload to persistent diabetic-like cardiac gene signatures.

In our study, the RNA sequencing of heart tissues of ISO/OP mouse models were conducted at 24h and 5 weeks post ISO exposure. The RNA sequencing identified ISO/OP DEGs associated with fatty acid metabolic process, carbon metabolism, and cardiomyocyte functions. One of the upregulated gene expressions is pyruvate dehydrogenase kinase 4 (Pdk4). Pdk4 is a key regulator of glucose metabolism in the heart and Pdk4 high expression interrupts cardiac glucose metabolism and associated with interruption of fatty acid metabolism [27, 53]. Another ISO upregulated gene, Serpine 1, has been known strongly associated with cardiovascular diseases such as atherosclerosis and heart failure [54, 55]. High expression of Serpine1 contributes to cardiomyocyte apoptosis in sepsis [56]. Furthermore, ISO exposure suppresses transferrin receptor (Tfrc) expression in the heart. Tfrc is responsible for iron transportation. It is critical for iron homeostasis for heart function. Lack of Tfrc causes iron deficiency and is associated with cardiomyopathy and heart failure [57]. In addition, ISO exposure may increase the risk of impaired cytoskeleton function by suppressing Tub8a gene expression. Tuba8 encodes an alpha-tubulin isoform expressed in heart tissue. It contributes to microtubule organization, structural integrity, and contractile function of cardiomyocytes. ISO exposure inhibits Tuba8 expression may lead to altered cardiomyocyte contractility.

When combined with prior studies on neurological deficits after GA/OP, our findings suggest that acute perioperative shifts in transcriptomic profiles may be an indicator for long-term cardiac vulnerability and a possible underlying mechanism for the development of brain dysfunction long after surgery in aged mice [15]. Studies demonstrated that chronic suppression of β-oxidation and lipid accumulation can trigger mitochondrial and ER stress, leading to insulin resistance and proteostatic failure—core features of diabetic CM [58, 59]. Similar metabolic and oxidative disruptions occur in the brain after anesthesia, driving synaptic remodeling and cognitive decline [60–62]. These shared transcriptomic patterns between heart and brain point to coupled dysregulation of fatty acid metabolism, redox balance, and ER stress. Mitochondrial dysfunction in the heart can release inflammatory mediators that modulate neuroinflammation [63–65], while neural mitochondrial injury may impair autonomic regulation of cardiac function, forming a bidirectional feedback loop [66–68].

The heart-brain axis has been acknowledged by many to be a contributing factor in neurodegenerative diseases, with many studies pointing to dysregulated heart function as a possible mechanism for Alzheimer’s and Parkinson’s disease [69–72]. Moreover, an abundance of evidence points to open-heart surgery leading to POCD [73–75], with one study even showing elevated levels of Abeta being detected in the cerebrospinal fluid of patients that underwent cardiopulmonary bypass [76]. However, only a few studies have examined the effects that GA may have on the cardiovascular system when patients don’t undergo heart operations [77]. Thus, we are one of the first groups to report on subtle molecular changes in cardiac tissue following ISO/OP. Importantly, our findings indicate high levels of ER stress being present in the heart at 24 h after surgery, a critical timepoint in the perioperative period. When combined with transcriptomic data of cardiac tissue taken from chronic 5-week animals, and the observations of neural deficits in our previous study, we can obtain a bigger picture of how the heart-brain axis mediates neurological consequences after ISO/OP in aged mice. Overall, the present study suggests that ER stress in the heart may be a potential mechanism and fascinating target for future studies on POCD and POD.

## Acknowledgements

This study was supported by the National Heart Lung and Blood Institute Grant R01 HL160727 (W-B Shen).

## Author Contributions

Y.L., J.G.L, and W.S. conceived the project and designed the experiments. W.W.Y. and A.W.C. contributed to conceptualization, hypothesis development, data analysis, and interpretation. Y.L. and H.Li. performed ISO/OP experiments and sample collection. J.G.L. prepared all tissue RNA extraction. H.Lee. performed RNAseq data analysis and prepared the figures. W.W.Y., A.W.C., Y.L., J.G.L., and W.S. wrote the manuscript. All authors approved the final version of the manuscript.

## Data Availability Statement

All data needed to evaluate the conclusions in the paper are present in the paper and/or the Supplementary Materials.

